# Neural patterns differentiate traumatic from sad autobiographical memories in PTSD

**DOI:** 10.1101/2022.07.30.502151

**Authors:** Ofer Perl, Or Duek, Kaustubh R. Kulkarni, Ben Kelmendi, Shelley Amen, Charles Gordon, John H. Krystal, Ifat Levy, Ilan Harpaz-Rotem, Daniela Schiller

## Abstract

For people with post-traumatic stress disorder (PTSD), recalling traumatic memories often displays as intrusions that differ profoundly from processing of ‘regular’ negative memories. These mnemonic features fueled theories speculating a qualitative divergence in cognitive state linked with traumatic memories. Yet to date, little empirical evidence supports this view. Here, we examined neural activity of PTSD patients who were listening to narratives depicting their own memories. An inter-subject representational similarity analysis of cross-subject semantic content and neural patterns revealed a differentiation in hippocampal representation by narrative type: Semantically similar sad autobiographical memories elicited similar neural representations across participants. By contrast, within the same individuals, semantically thematically similar trauma memories were not represented similarly. Furthermore, we were able to decode memory type from hippocampal multivoxel patterns. Finally, individual symptom severity modulated semantic representation of the traumatic narratives in the posterior cingulate cortex. Taken together, these findings suggest that traumatic memories are a qualitatively divergent cognitive entity.

## Main

The involuntary re-experiencing of the traumatic autobiographical memory, often following exposure to trauma-related stimuli, is a hallmark feature of post-traumatic stress disorder (PTSD)^1, 2^. Although personal memory is at the core of PTSD symptoms, research on the neural mechanism of PTSD has largely focused on non-personal basic learning and memory paradigms^3^. It is yet unclear whether traumatic memories differ from negative non-traumatic autobiographical memories in a qualitative or a quantitative manner. Is a traumatic memory an exceptionally strong manifestation of autobiographical memory or a divergent neural representation of memory?

To examine this, we need to factor-in differences across individual traumatic narratives and the idiosyncratic experiences they evoke, and extract from them the common markers operating in trauma-driven state. With this in mind, we designed a study that examines PTSD patients’ neural responses to their own personal traumatic memory in the form of a structured, fully annotated, audio narrative. We compared traumatic memory, within each participant, to a negatively-valenced non-traumatic sad memory, and a calm positive memory.

Previous research has widely established the role of the hippocampus in the construction of relational cognitive maps, onto which events are bound across space and time to form episodic memories^4–6^. It is through this tracking of sequences of events that the hippocampus generates a narrative from discrete events^7, 8^. In turn, the hippocampus also governs the ensuing retrieval of such events^9, 10^. The hippocampus is in fact so central to maintenance of episodic memory that lesioning it results in grave deficits in mnemonic abilities, to the point of global anterograde amnesia in humans^11^.

Impairments to hippocampal processes are at the heart of PTSD pathophysiology^12^. A solid body of evidence suggests that PTSD is associated with structural abnormalities (predominantly a reduction in volume), as well as reduced functional connectivity between the hippocampus and other regions of the default mode network during rest^13, 14^. In the context of encoding of the traumatic memory itself, peri-traumatic aberrations in hippocampal functions are thought to contribute to the paradoxical mnemonic sequalae commonly observed in PTSD – difficulty in voluntary coherent recall alongside with detailed involuntary intrusions of the traumatic memory^15, 16^. All in all, our understanding of the impact of PTSD on the spectrum of hippocampus-mediated mnemonic processes, is still murky^17^

Emotional memories—episodic memories that elicit emotions at retrieval—often engage the amygdala^18^. Functionally, the amygdala is considered one component in a broader neurocircuitry model implicated in PTSD, comprised of interactions with the hippocampus and medial prefrontal cortex^19^. Generally, the amygdala shows hyperresponsivity to both non-specific threat-related stimuli^20^ and personal trauma reminders in PTSD (Rauch et al., 1996; Shin et al., 2004; also see meta-analysis by Etkin and Wager, 2007). Amygdala-hippocampus connectivity during construction of negative autobiographical memories has also resulted in conflicting findings demonstrating decreased^3, 24^, but also increased functional connectivity^25^ between these two structures.

Amygdala activation during memory encoding modulates the memory’s explicit subsequent strength^26^, evaluated through its persistence, accuracy, and vividness^27^. Amygdala activity can also be driven by hippocampal inputs during the reinstatement of an aversive memory. Such hippocampal input is typically based on episodic retrieval, however, semantic information may also induce fear (e.g., ‘Cobra snakes are dangerous’). In fact, amygdala responses may even arise from imagined stimuli, as in the case of experiences conveyed through narrative (e.g., horror or suspense fiction)^28^. Whether the amygdala itself serves as an ‘auxiliary’ site to support the storage of emotional memory is still debated^29–31^.

Considering this evidence, we examined whether and how the hippocampus and amygdala differentiate between traumatic and sad autobiographical memories. We hypothesized that across PTSD patients, semantic similarity would correspond to neural similarity: if the personal memories of two participants are semantically close, their patterns of neural responses while listening to the audio recording of these memories should be similar as well. If traumatic and sad memories are just different cases of autobiographical memories, we should observe semantic-to-neural correspondence across pairs of traumatic memories and pairs of sad memories alike. However, if traumatic autobiographical memories diverge from—rather than being a version of—sad autobiographical memories, then we would observe the semantic-to-neural relationship only for sad, but not traumatic memories. Our hypothesis further suggests that a shared neural representation will allude to a shared underlying neural mechanism. If the effect does not extend to pairs of traumatic memories across participants, despite their semantic similarity, this may imply traumatic memories diverge from the neurotypical mechanisms of other sad, non-traumatic, autobiographical memory.

We hypothesized that if such mechanistic difference between traumatic and sad memories exists, it would be detected in hippocampal neural patters. In contrast, and as a control comparison, we did not expect such differentiation, or any pattern representation related to the semantic content of memories in the amygdala, given the role of this region primarily in signifying emotional valence.

## Results

Twenty-eight participants (age = 38.2 ± 10.4 years, 11 females) diagnosed with PTSD (CAPS score = 41.2 ± 8.3), underwent reactivation of autobiographical memory through script-driven imagery while undergoing functional magnetic resonance imaging (fMRI). First, in order to generate stimuli which are based on participants’ autobiographical memory, we used an imagery development procedure. Participants elaborated on three types of autobiographical memories: 1) the ‘PTSD’ condition: the traumatic event associated with their PTSD (DSM-5 criterion A; common examples were combat, sexual assault, domestic violence), 2) the ‘Sad’ condition: a sad meaningful, but non-traumatizing experience (common examples were death of family member or pet) and 3) the ‘Calm’ condition: a positive, calm event (common examples were memorable outdoor activities). These highly personal and variable depictions of autobiographical memory were then systematically arranged into an approximately 120-second audio clip (referred to henceforward as ‘script’ or ‘narrative’, interchangeably), narrated by a member of the research staff. All scripts were composed with ample attention to a common rigid structure, into which the individual autobiographical memory was incorporated. Notably, ‘PTSD’ and ‘Sad’ narratives were scripted to maximize their structural similarity to control for content and arousal (see Methods). Participants listened to this novel rendition of their autobiographical memory for the first time while undergoing functional magnetic resonance imaging (fMRI) (**Figure 1A**).

**Figure 1:**
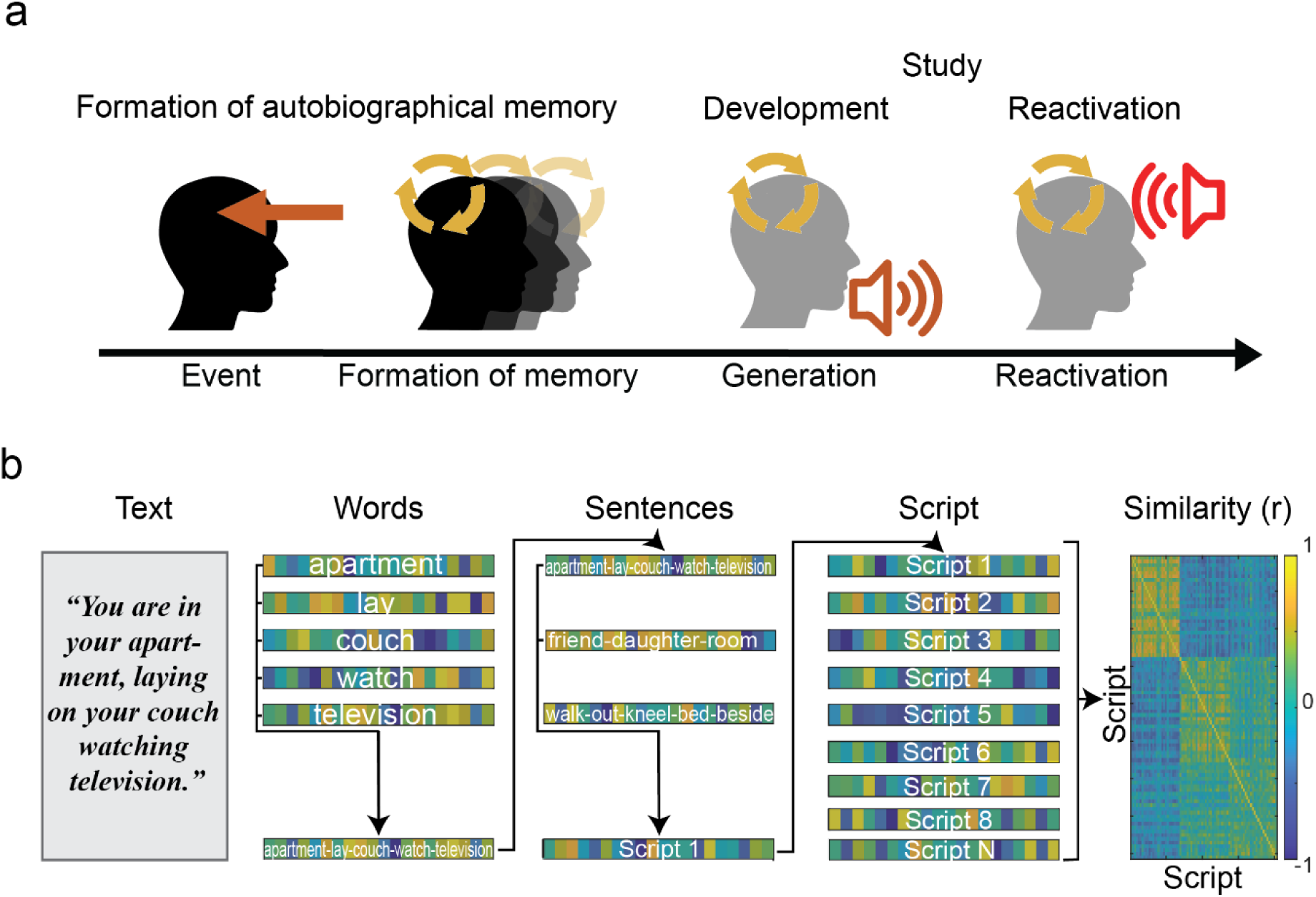
**(a) Experimental paradigm for script-based autobiographical memories reactivation.** At some point prior to enrollment, an event was perceived, and an autobiographical memory was formed. This memory has since then undergone an unknown number of recalls and reconsolidation iterations. During the study, participants once again recalled this memory, this time while verbalizing it as part of the imagery development procedure. These recollections were incorporated into a novel narrative-form rendition of the autobiographical memory, which was played to the participants for the first time while undergoing functional imaging, in order to reactivate this autobiographical memory once again. ***(b)* Semantic similarity of autobiographical narratives using word embedding.** Each indexable word in the script was assigned a 300-dimensional vector representation (e.g., ‘apartment’, ‘lay’ etc.). Sentence vectors were represented as the average of word vectors comprising them (e.g., tokenized e.g., ‘apartment lay couch watch television’) and scripts represented as the average of the sentences in them. Pairwise semantic similarity across participants was calculated.

Basic group inference in cognitive neuroscience, and neuroimaging in particular, relies on the detection of shared stimulus-induced signals of greater amplitude than the noise or idiosyncratic signals in these systems^32^. Autobiographical memories are rarely generated in a controllable lab setting, and therefore differ in their content. However, at the same time, autobiographical memories recall may elicit common cognitive states (e.g., mental time travel) that are potentially subserved by common neural substrates across individuals. In this study, the sensory stimuli used for reactivation were based on idiosyncratic experiences in order to invoke a common cognitive state – the reexperiencing of a traumatic autobiographical memories. During recruitment participants were not screened for a specific trauma type. This enabled us to span a wide range of themes, some of which were present in ‘PTSD’ and ‘Sad’ conditions. For example, a narrative describing the death of a loved one can meet PTSD criterion A classification for one participant and thus be associated with a traumatic autobiographical narrative yet be regarded as ‘Sad’ autobiographical memory (i.e., non-traumatizing) for another. This granted the opportunity to measure similarity of autobiographical memories in a parametric and continuous manner, and critically – to compare the representation of relatable autobiographical memories in light of their clinical outcomes – ‘PTSD’ or non-traumatizing.

### Semantic analysis of similarity in autobiographical memory

To quantify similarity between autobiographical memory based narratives across individuals and conditions we applied a word embedding approach – a computational linguistic tool used to quantify distances between text-based semantics^33^. In brief, words are pre-sampled from gigantic text corpora and are then embedded in a high-dimensionality space according to local co-occurrences. The derived semantic space allows to infer relational structure between concepts according to their distance. Such tools were previously used to uncover neural representations of semantic spaces^34^ both with functional imaging^35^ and invasive recordings^36^.

We used MATLAB’s word2vec with a pre-trained embedded space for one million words in the English language^37^. Each word was assigned a 300-dimensional vector representation. In our analytical hierarchy, sentence vectors were represented as the average of word vectors comprising them. Similarly, scripts represent as the average representation of their sentences (**Figure 1B**). The high dimensionality of the semantic dataset is difficult to interpret visually. We therefore applied *t-*distributed Stochastic Neighbor Embedding (*t*-SNE), a method for dimensionality reduction, to the data to cluster narratives based on the semantic similarity of their content and projected this dimensionality-reduced dataset onto a three-dimensional space.

We observed that both types of negatively-valenced narratives – ‘PTSD’ and ‘Sad’ – formed overlapping clusters in semantic space, whereas ‘Calm’ narratives were grouped in a separate part of the space (**Figure 2A**). Additional 2D projections of the semantic space are available in (**Figure S1**). This qualitative visualization affirmed that semantic content of ‘PTSD’ and ‘Sad’ autobiographical memories are comparable and thus ‘Sad’ scripts are poised to provide a valid control for the ‘PTSD’ scripts.

**Figure 2:**
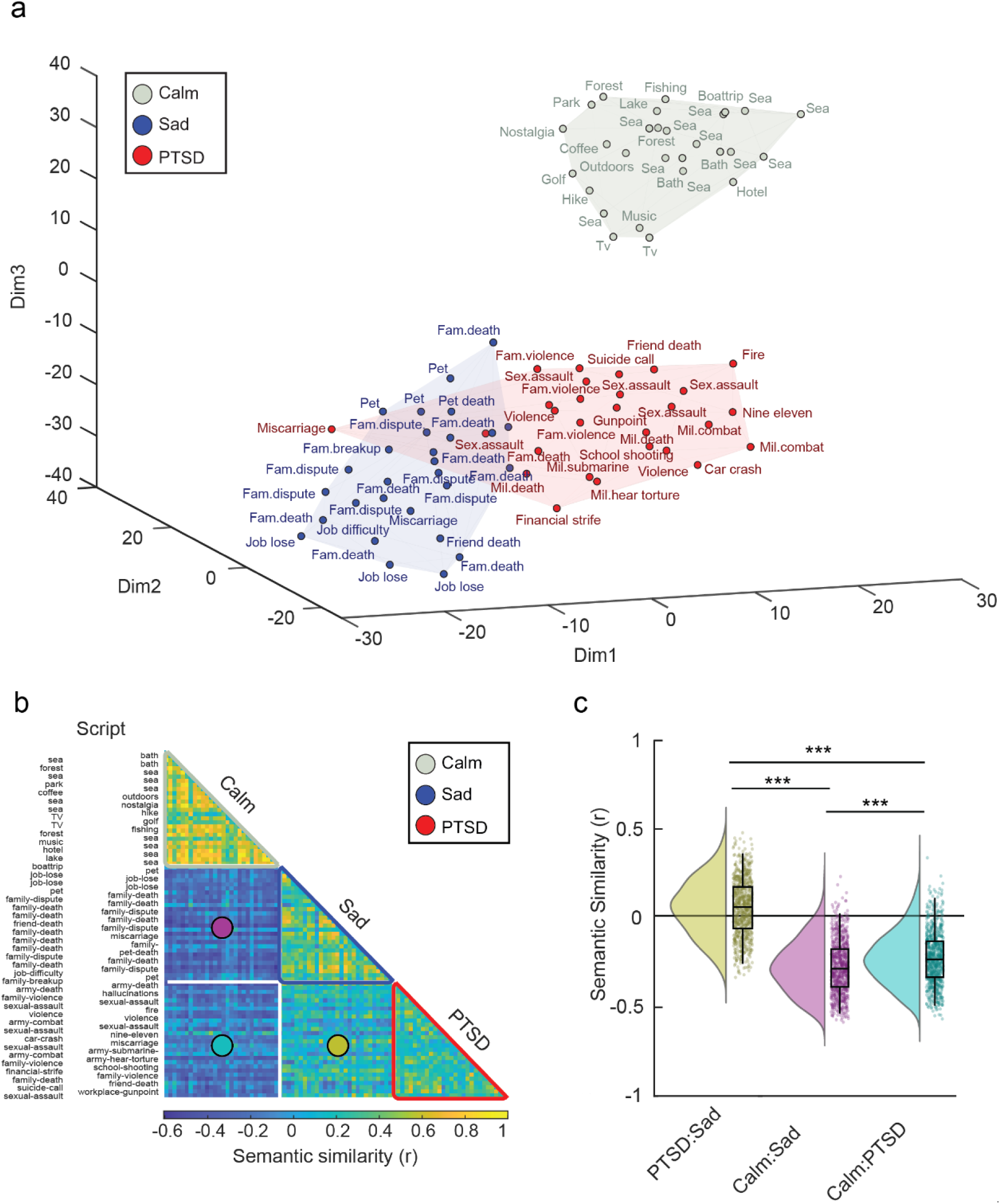
**(a) Clustering of semantic similarity across script types using t-SNE.** t-SNE embedding of scripts. Each dot represents a single script, projected onto a 3D space. Colored volumes are continuous areas in space occupied by each script type. Color denotes script type (‘PTSD’: red, ‘Sad’: blue, ‘Calm’: gray). Note overlap of ‘PTSD’ and ‘Sad’ semantic content. Text adjacent to data points are general titles of narrative content, generated by the researchers (abbreviations: Fam.: family, Sex. sexual. Mil. military). Some titles were omitted to prevent clutter. ***(b)* Pairwise similarity of semantic content of scripted narratives.** Semantic pairwise similarity (Pearson’s correlation coefficient, r) presented for all scripted narratives. Within-category similarity is marked in colored triangles off the matrix diagonal (‘PTSD’: red, ‘Sad’: blue, ‘Calm’: gray). Across-category similarity is marked in colored circles in distinct square sectors of the matrix (‘PTSD:Sad’: yellow, ‘Calm:Sad’: magenta, ‘Calm:PTSD’: teal). Text labels next to matrix rows are general titles of narrative content, generated by the researchers. ***(c)* Semantic similarity across script types.** Raincloud plots illustrating the distribution of pairwise semantic similarity between pairs of script types (‘PTSD:Sad’: yellow, ‘Calm:Sad’: magenta, ‘Calm:PTSD’: teal). *** p < 0.001

We next measured semantic similarity using the cosine distance between the 300-dimension vectors representing the scripts. The resulting semantic similarity matrix was comprised of the three script types (‘PTSD’, ‘Sad’, ‘Calm’) of 28 participants yielding a 84 X 84 matrix in total. Script types were grouped to aid in visualization of ‘type’ clusters off the diagonal (**Figure 2B**).

In addition to dimensionality reduction, we calculated cross-category similarity and observed that the ‘PTSD’ and ‘Sad’ narratives showed a higher cross-category semantic similarity than other cross-category comparisons (Similarity (r): ‘PTSD’:’Sad’ = 0.059 ± 0.159, ‘Calm’:’Sad’ = -0.274 ± 0.147 ‘Calm’:’PTSD’ = -0.224 ± 0.150; ANOVA: F(2,2351) = 1094.75, p < 1E-300. η_p_^2^ = 0.325) **(Figure 2C**). This semantic resemblance can be attributed to the themes shared by both negatively valenced scripts, as well as the systematically controlled structure and the shared phrases that were used.

In a complementary analysis we quantified pairwise similarity within each of the three script categories. As expected, we observed that within-category scripts displayed high semantic similarity (Similarity (r): ‘PTSD’ = 0.164 ± 0.139, ‘Sad’ = 0.237 ± 0.174 ‘Calm’ = 0.413 ± 0.174; 1-sample *t-*test vs 0:, all p < 1E-300). Further, an ANOVA conducted on script types revealed a main effect of ‘script’ (ANOVA: F(2,1133) = 231.80, p = 4.29E-85, η_p_ ^2^ = 0.225). Post-hoc *t-*tests further demonstrated that ‘Calm’ scripts had higher within-category similarity than ‘Sad’, and ‘Sad’ scripts had higher with-category similarity than ‘PTSD’ (both p < 0.0001, corrected with Tukey’s HSD).

These results suggest that sad and traumatic memories in the cohort overlapped in themes and semantic content. This analysis laid foundations for asking whether the neural patterns associated with these memories will differ by their clinical classification - traumatic or sad. Thus, any ensuing neural differences between ‘PTSD’ and ‘Sad’ reactivations may present themselves on top of a maximally identical pool of stimuli. We note that given their autobiographical nature and the use of a naturalistic paradigm, such stimuli may never be identical. However, establishing a “handle” on the differences and commonalities of the narratives enables us to leverage those differences in a quantifiable manner to provide neural insight.

### Intersubject Representational Similarity Analysis (IS-RSA)

All participants listened to each script type three times during the functional scan. Given our a-priori interest in the involvement of the hippocampus and amygdala in the processing of traumatic autobiographical memories in PTSD, we extracted signals from these structures using the Harvard-Oxford probabilistic atlas to conduct region-of-interest (ROI) analyses. In order to enhance signal to noise for the detection of the common pattern elicited by the repeated stimuli, the time courses were averaged across the three repeats of the script per ROI. The voxel time course of this average run was collapsed across time to generate a spatial pattern associated with the reactivation of each autobiographical memory. Finally, we generated a neural similarity matrix by calculating the pairwise Pearson correlation coefficient between the spatial patterns of all scripts. The dimensions of this matrix were identical to the one storing the narratives’ semantic similarity (28 participants X 3 script types) (**Figure 3A**). To account for the idiosyncratic nature of autobiographical memories, we used intersubject representational similarity analysis (IS-RSA) to relate the personalized semantic content of the scripts with neural representations acquired during script reactivation. In IS-RSA, the neural and semantic similarity matrices were vectorized and correlated (Spearman’s correlation) within each category of script type, yielding three correlation coefficients (‘rho’, ρ) per ROI, one for each script type (**Figure 3B**). The first IS-RSA we conducted related between-participant neural similarity of hippocampal patterns (**Figure 3C**) with the between-participant semantic similarity matrix computed before (**Figure 3D**).

**Figure 3:**
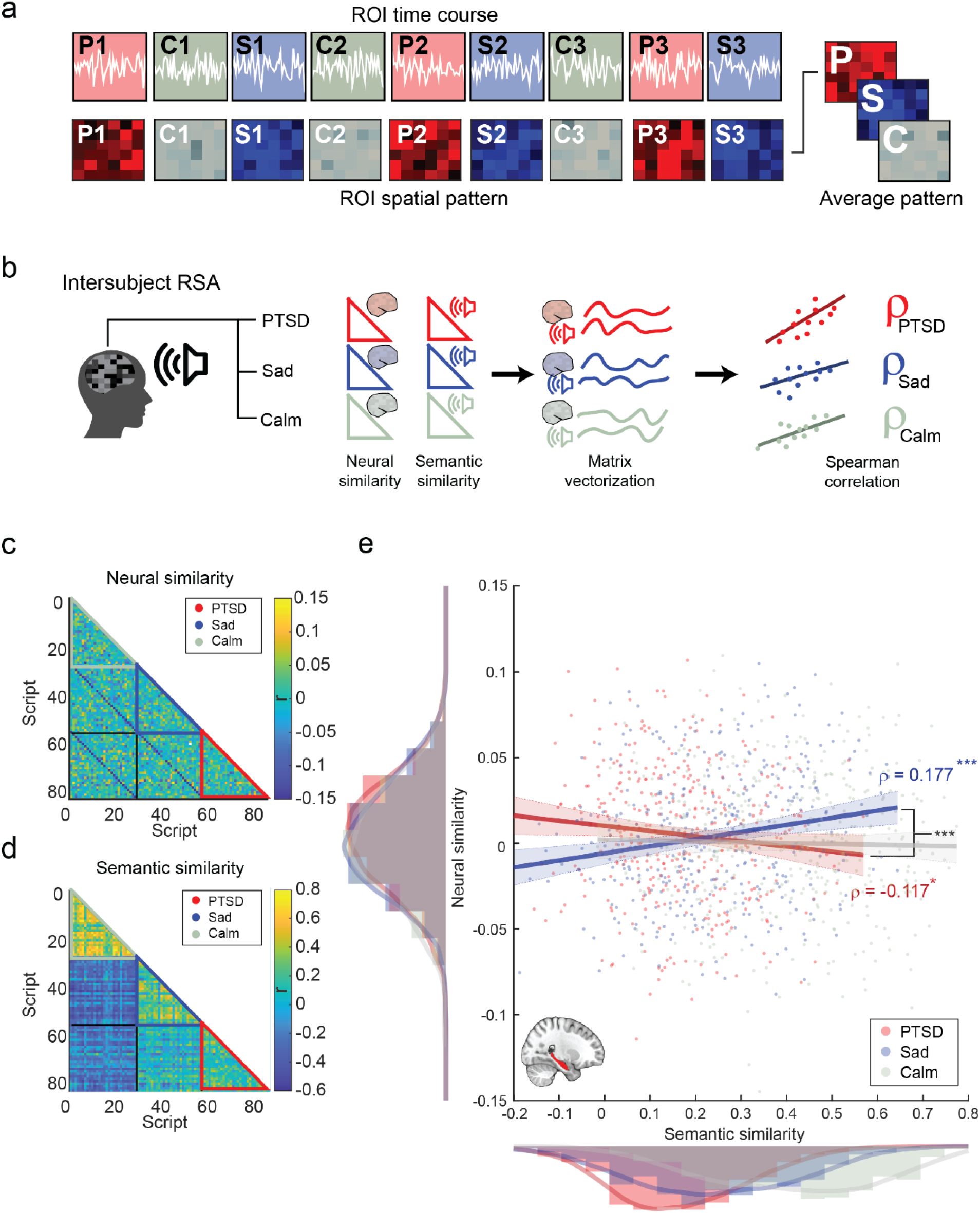
**(a) Extraction of spatial patterns.** Each script narrative (‘PTSD’, ‘Sad’, ‘Calm’) was played three times in the scanner. spatial patterns were extracted from regions of interest and averaged across repeated presentations to generate an average pattern associated with each script reactivation. *(b)* **Intersubject representational similarity analysis (IS-RSA).** We carried out three independent IS-RSA in which intersubject variability during script reactivation was captured using two subject by subject matrices: one depicting neural pattern pairwise similarity, and the other depicting semantic pairwise similarity. Spearman rank correlation was calculated for each pair of vectorized similarity matrices to provide a correlation coefficient tying semantic and neural representation of either ‘PTSD’, ‘Sad’ or ‘Calm’ narratives. *(c)* **Hippocampus – neural similarity matrix.** Script-by-script neural similarity matrix for spatial patterns extracted from the hippocampus during script reactivation. Within-category similarity is marked in colored triangles off the matrix diagonal (‘PTSD’: red, ‘Sad’: blue, ‘Calm’: gray). *(d)* **Semantic similarity matrix** Script-by-script semantic similarity of scripted narratives. Within-category similarity is marked in colored triangles off the matrix diagonal (‘PTSD’: red, ‘Sad’: blue, ‘Calm’: gray). *(e)* **Hippocampus – semantic-to-neural IS-RSA.** Intersubject representational similarity analysis conducted on pairwise similarity of semantic content and neural patterns in the hippocampus. Each datapoint is one pairwise comparison. Analysis was iterated per script type (‘PTSD’: red, ‘Sad’: blue, ‘Calm’: gray). Histograms along axes depict similarity distribution, thick trace depict estimated density, colors correspond to main legend. Regression lines are approximate visualization of Spearman correlation rho coefficients for IS-RSA in ‘PTSD’ and ‘Sad’ scripts (red and blue resp.). * p < 0.05; *** p < 0.001.

In the hippocampus, the IS-RSA uncovered a differentiation in semantic representation: semantic similarity scaled positively with neural similarity for ‘Sad’ narratives (‘Sad’, ρ = 0.177, p_(FDR corrected)_ = 0.0034) but not for ‘PTSD’ narratives (‘PTSD’, ρ = -0.117, p_(FDR corrected)_ = 0.069) (**Figure 3E**). (We further verified the strength of link between semantic and neural matrices in the ‘Sad’ condition using Mantel’s test for matrices correlation^38, 39^ which suggested a strong effect, p = 0.001).

We then tested whether the two correlation coefficients of IS-RSA conducted on ‘PTSD’ and ‘Sad’ scripts, and associated hippocampal patterns differed significantly. The correlation coefficients underwent a z-score transformation and the absolute difference between them was assigned with a p value (se Methods). We observed that indeed hippocampal IS-RSA representations of semantic content significantly differed as a function of script type (Coefficient comparison (two-tailed) PTSD’ vs. ‘Sad’, hippocampus: p_(FDR corrected)_ = 0.0003.

### Control Analyses

We verified that the difference in semantic-to-neural mapping is not due to differences in mean amplitude between the conditions. To this end we conducted general linear modeling (GLM) on the functional imaging data in which regressors related to all event types were created. The GLM included three separate regressors denoting the full duration of the three types of scripts – ‘PTSD’, ‘Sad’ and ‘Calm’. We extracted univariate parameter coefficients from the hippocampus and compared them using a repeated-measures ANOVA (rmANOVA) which uncovered no differences between script types (rmANOVA F(2,27) = 0.67, p = 0.528).

To control for possible habituation over the course of script playback, we repeated this approach while modeling the early and late parts of the script separately (roughly one minute each). Focusing on the early part, we again observed comparable parameter estimates across script types (rmANOVA F(2,27) = 1.59, p = 0.212). Taken together, these results mitigate concerns regarding confounding effects of different arousal levels across conditions.

We also sought to rule out a possible confounding factor in narrative similarity that may have been driven by similarity of low-level acoustic properties of the auditory stimuli. To this end we generated an acoustic similarity matrix (see Methods) and conducted an IS-RSA using the acoustic and neural similarity matrices. We did not observe any significant representation of acoustic features in the hippocampal patterns, both in ‘PTSD’ and ‘Sad’ conditions (‘PTSD’: ρ = 0.042 p = 0.41, ‘Sad’: ρ = -0.037, p = 0.48) (**Figure S2**).

### Investigation of hippocampal subregions involvement in representation

Evidence suggests hippocampal subregions along its longitudinal axis are recruited differently during tasks involving recall autobiographical memory and scene construction^40, 41^ with particular implications in PTSD^42–44^. With this in mind we conducted an exploratory analysis to further delineate the differences in patterns of hippocampal representation of traumatic autobiographical memory narratives. We conducted IS-RSA on neural patterns extracted separately from the anterior and posterior extremities of the hippocampus (split into three segments: anterior, middle, and posterior, see Methods).

We observed the semantic representations were more pronounced in the posterior part of the hippocampus (Posterior hippocampus: ‘PTSD’, ρ = -0.0506, p = 0.327; ‘Sad’, ρ = 0.169, p_(FDR-corrected)_ = 0.0004; Coefficient comparison (two-tailed) ‘PTSD’ vs. ‘Sad’ p_(FDR-corrected)_ = 0.0005 ; anterior hippocampus: ‘PTSD’, ρ = -0.0503, p = 0.329; ‘Sad’, ρ = 0.065, p = 0.204; Coefficient comparison (two-tailed) ‘PTSD’ vs. ‘Sad’ p = 0.112) (**Figure S3**).

### No evidence for semantic-to-neural patterns in the amygdala

To examine whether the lack of semantic representation of traumatic autobiographical memories was specific to the hippocampus, we repeated the same IS-RSA with neural patterns extracted from the amygdala. Signals from the amygdala did not demonstrate a significant link between semantic content and neural patterns for either ‘Sad’ (ρ = 0.066, p_(FDR corrected)_ = 0.323) or ‘PTSD’ narratives (ρ = -0.057, p_(FDR corrected)_ = 0.323) (**Figure S4)**. In addition, a univariate analysis of GLM-derived parameter estimates suggested comparable levels of activation across script types (rmANOVA F(2,27) = 1.21, p = 0.307).

We verified that this null effect is not a result of averaging left and right amygdala signals. We conducted IS-RSA and univariate analysis left and right amygdala separately and observed no differences between ‘PTSD’ and ‘Sad’ conditions (IS-RSA: Left amygdala: ‘PTSD’, ρ = -0.03, p = 0.56, ‘Sad’, ρ = 0.024, p = 0.643; Coefficient comparison (two-tailed) ‘PTSD’ vs. ‘Sad’ p = 0.46); Right amygdala: ‘PTSD’, ρ = -0.064, p = 0.21, ‘Sad’, ρ = 0.078, p = 0.129; Coefficient comparison (two-tailed) ‘PTSD’ vs. ‘Sad’ p = 0.051); parameter estimates: Left amygdala (F(2,27) = 1.36, p = 0.265; Right amygdala (F(2,27) = 0.78, p = 0.463).

### Autobiographical memory type can be decoded from hippocampal patterns

To further validate that spatial patterns in the hippocampus, but not the amygdala, carry meaningful information about the condition associated with these negative autobiographical memories, we trained a regularized linear discriminant analysis (rLDA) model to decode narrative conditions (‘PTSD’ or ‘Sad’) from the multivoxel spatial patterns extracted from the hippocampus during script reactivation. Using 25-fold cross-validation we were able to decode at 66.2 % accuracy whether hippocampal spatial patterns belonged to a ‘PTSD’ or ‘Sad’ narrative. To assess the power of this predictive ability, we iterated the procedure with shuffled labels (N = 2,500) to obtain a surrogate distribution of decoding accuracy (mean ± SD = 52.7 % ± 4.9). Nonparametric testing confirmed that decoding accuracy for the empirical data was well above chance (p = 0.0028) (**Figure 4A**). To further validate our finding, we then attempted to decode ‘PTSD’ from ‘Calm’ scripts, expecting those to be teased apart more easily given their different mental state. Indeed, we were able to decode these conditions at 80.9 % accuracy, which was well above chance (mean ± SD = 54.2 % ± 5.0, p < 0.0001) (**Figure 4B**). For comparison, spatial patterns derived from the amygdala could not be used as robustly to distinguish between ‘PTSD’ and ‘Sad’ scripts (decoding accuracy for empirical data: 58.2 %, surrogate: mean ± SD = 52.2 % ± 4.7, p = 0.097) (**Figure 4C-D**). These findings provide additional support to the idea that hippocampal activity during traumatic autobiographical memory recall represents elements of the narrative, and these patterns are observably different from activity during non-traumatizing autobiographical memory recall.

**Figure 4:**
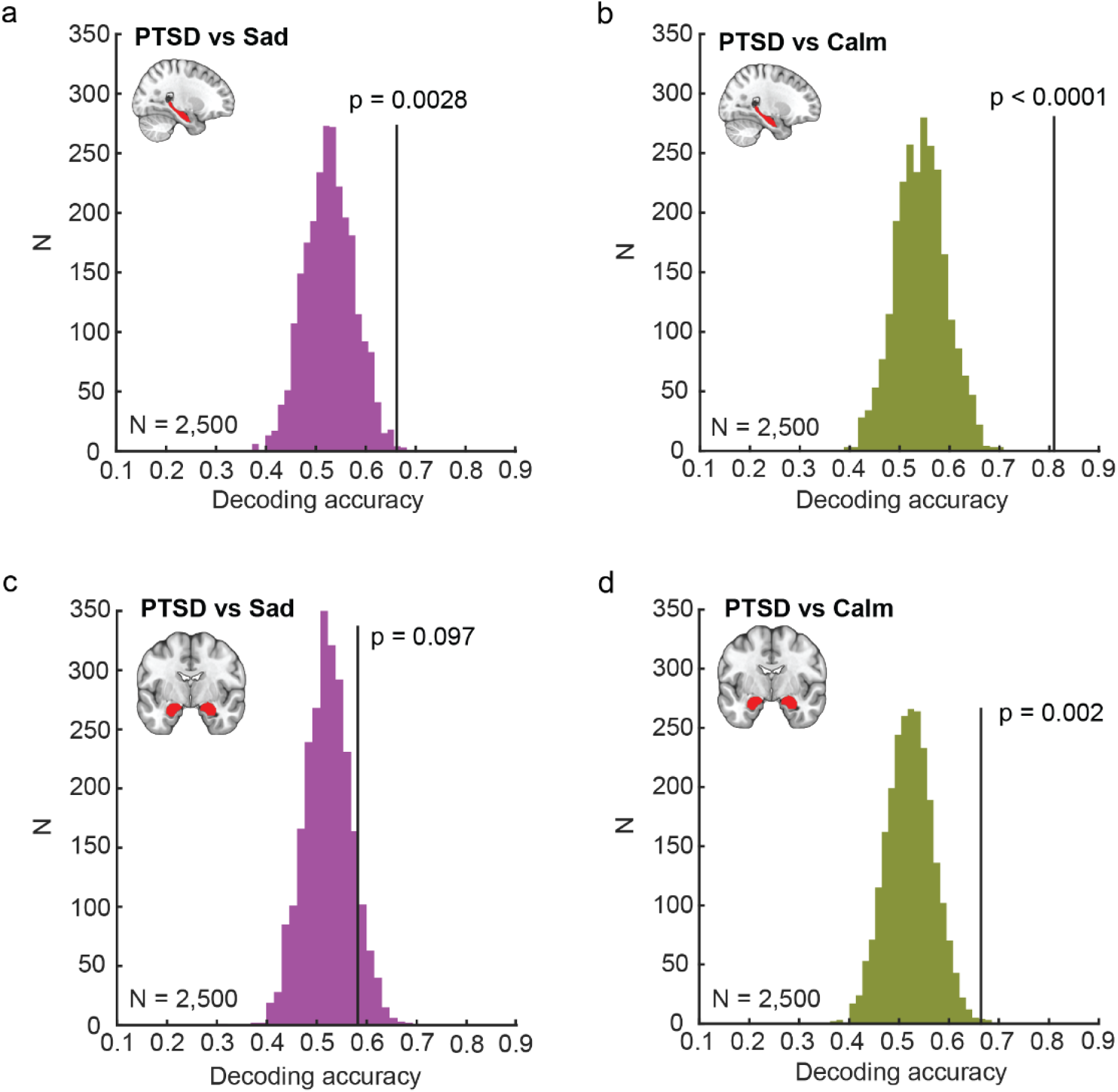
**(a) Decoding accuracy of scripted narrative type from neural patterns.** Vertical black line denotes decoding accuracy of ground truth script type (‘PTSD’ vs. ‘Sad’) from hippocampal spatial patterns in the empirical condition. Colored histogram is a surrogate distribution comprised of decoding accuracy for the same neural data with shuffled labels. p value is derived non-parametrically through a permutation test (N = 2,500). (b) Same as **(a)** but for decoding accuracy of ‘PTSD’ vs. ‘Calm’ from hippocampal spatial patterns. (c) Same as **(a)** but for decoding accuracy of ‘PTSD’ vs. ‘Sad’ from amygdala spatial patterns. (d) Same as **(a)** but for decoding accuracy of ‘PTSD’ vs. ‘Calm’ from amygdala spatial patterns.

### Semantic representation for the ‘PTSD’ narratives in the posterior cingulate cortex

Having observed that hippocampal representations for narratives of traumatic autobiographical memory are diminished, compared to non-traumatizing events, we expanded our investigation to ask whether any brain regions will show better semantic representation for the ‘PTSD’ narratives over ‘Sad’ ones. We hypothesized that the posterior cingulate cortex (PCC), a hub of the default mode network known to be heavily implicated in both narrative comprehensive and autobiographical memory processing^32, 45^, and particularly in emotional memory imagery ^46^, will be a candidate region sensitive to script reactivation.

To define the PCC functionally, we generated a contrast comparing neural activity during narrative playback (all three script types were included) to inter-trial interval baseline. This contrast therefore factored activations for both positive and negative valence scripts. When we applied IS-RSA to neural patterns extracted from the PCC functional ROI we found an overall lack of semantic representation in that ROI (‘PTSD, ρ = 0.077, p_(uncorrected)_ = 0.136, ‘SAD, ρ = 0.017, p_(uncorrected)_ = 0.745).

Since our patient cohort was heterogenous in terms of symptoms severity and background, we asked whether PTSD severity, operationalized as CAPS-5 score evaluated during screening, will explain the between-subject variability in the extent by which narratives will be represented in the PCC. To this end, we split the cohort into two sub-groups (both N = 14) labelled ‘High’ and ‘Low’ according to their CAPS-5 score. The total scores in the study ranged between 26 – 60 (mean ± SD = 41.2 ± 8.3). Setting a cutoff CAPS score of 38 (inclusive), the two resulting sub-groups significantly differed in their CAPS score (‘High’ = 47.8 ± 3.4, ‘Low’ = 34.7 ± 6.5 t(26) = 6.68, p < 0.0001).

First, we verified that univariate BOLD responses in these ROIs were comparable. We used a whole-brain two-sample *t-*test design to compare the ‘High’ and ‘Low’ symptoms groups’ response across script types. In both ‘PTSD’ > ‘Calm’ and ‘PTSD’ > ‘Sad’ contrasts we were not able to detect differences between the groups using cluster size threshold corrected with p_FWE_ < 0.05. Next, we verified that semantic similarity within these sub-groups was comparable, that is, that script similarity within each script category did not differ between the groups. In ‘PTSD’ scripts, the average semantic similarity did not differ between the ‘High’ and ‘Low’ symptoms groups (two-sample *t-*test, t(180) = -0.73, p = 0.46); ‘Sad’: t(180) = -2.03, p = 0.043 (non-significant after Bonferroni correction for multiple comparisons 0.05 / 3 = 0.0167)). In ‘Calm’ condition, scripts similarity was significantly higher in the ‘Low’ group than the ‘High’ one (‘Calm’: t(180) = -3.18, p = 0.0017. We therefore applied ensuing IS-RSA only to the ‘PTSD’ and ‘Sad’ conditions.

We conducted IS-RSA separately for each subgroup, using its corresponding semantic and neural similarity matrices. Matrix dimensions were 14 X 14, which when vectorized, resulted in 91 values. We compared the resulting Spearman coefficient that were derived for each of the two symptom groups using a nonparametric test. We split the full cohort into two random sub-groups iteratively (N = 25,000) and computed a surrogate distribution to which the statistics of the true, CAPS-5-based split, were compared.

The PCC displayed a strong discriminative utility where higher symptom severity was associated with stronger semantic representation of ‘PTSD’ scripts (**Figure 5A**). Representation of the ‘Sad’ scripts also showed stronger representation in the ‘High’ symptoms group, however to a much lesser extent (**Figure 5B**). (PCC ‘PTSD’: ‘High’: ρ = 0.266; ‘Low: ρ = 0.0687; ‘Sad’: ‘High’: ρ = 0.0756; ‘Low: ρ = -0.0537). By contrast, in the hippocampus, the extent of the link between neural patterns and semantic representations of both ‘PTSD’ and ‘Sad’ scripts did not differ by symptom severity (hippocampus ‘PTSD’: ‘High’: ρ = -0.254; ‘Low: ρ = -0.163; ‘Sad’: ‘High’: ρ = 0.199; ‘Low: ρ = 0.206). Nonparametric permutation tests asserted the Spearman coefficients significantly differed by symptom severity in the IS-RSA based on PCC, but not the hippocampal neural patterns (PCC coefficient difference_(High - Low)_ = 0.197, nonparametric permutations p = 0.0001; ‘Sad’: coefficient difference_(High - Low)_ = 0.129, nonparametric permutations p = 0.0036; hippocampus ‘PTSD’: coefficient difference_(High - Low)_ = -0.074, nonparametric permutations p = 0.729; ‘Sad’: coefficient difference_(High - Low)_ = -0.007, nonparametric permutations p = 0.885) (**Figure 5C**). A summary of IS-RSA correlation coefficients per ROI and symptoms group conveys a differentiation by symptoms that is present in the PCC signals but is almost entirely absent in the hippocampus (**Figure 5D**). This result suggests that the severity of PTSD symptoms is linked to the semantic representation of the traumatic narrative in the PCC, whereas the differentiation in representation observed in the hippocampus persisted regardless of symptoms severity.

**Figure 5:**
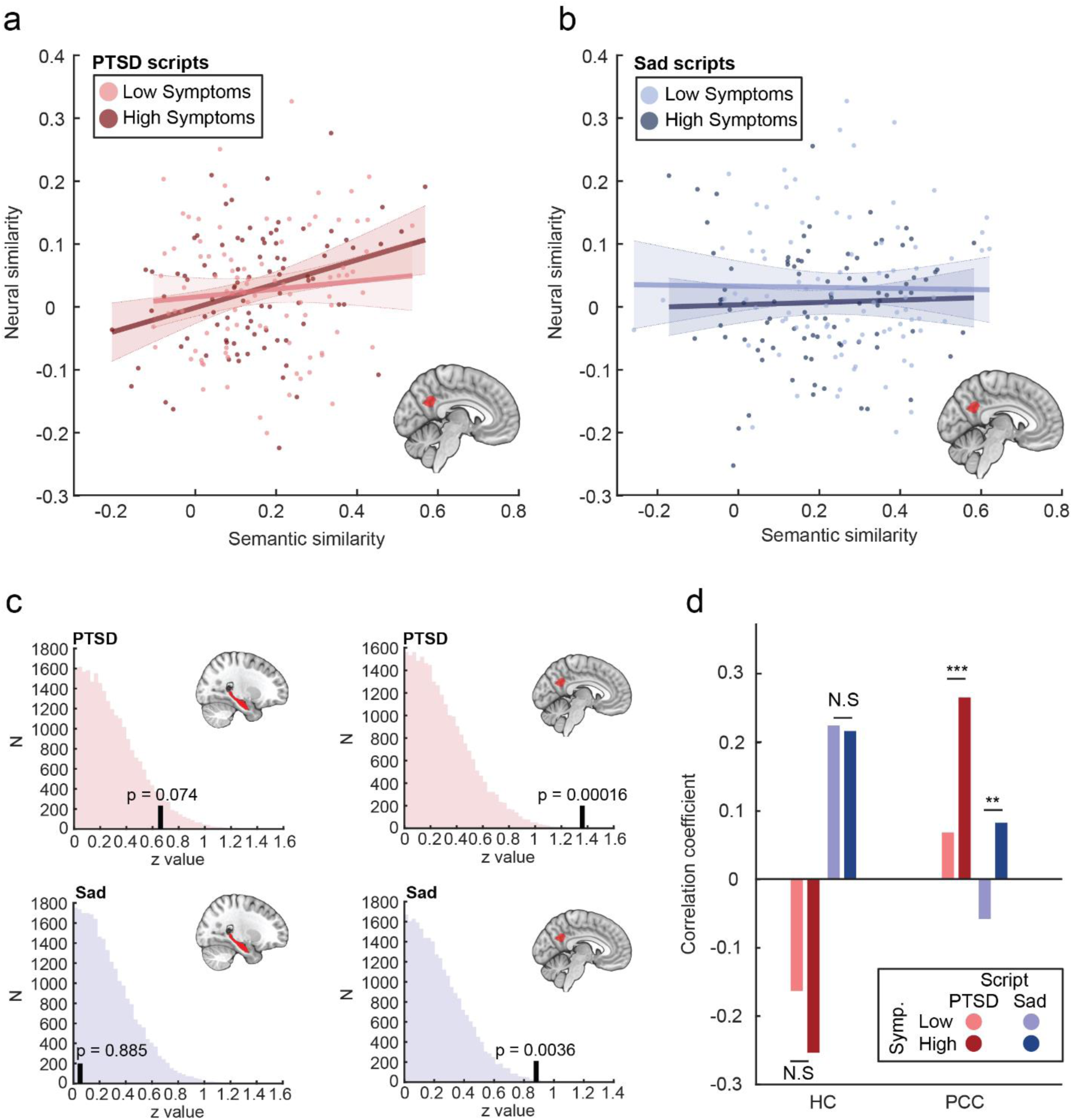
**(a) Intersubject representational similarity analysis in the PCC differed by symptom severity** IS-RSA conducted on pairwise similarity of semantic content of ‘PTSD’ narratives and neural patterns in the PCC on subgroups differing in symptom severity (‘Low’ and ‘High’). Each datapoint is derived from a pairwise comparison. Analysis was iterated per subgroup. Regression lines are approximate visualization of Spearman correlation rho coefficients for IS-RSA in ‘Low’ and ‘High symptoms (light and dark red, respectively.). (b) Same as **(a)** but conducted on semantic content of ‘Sad’ narratives and neural patterns in the PCC on subgroups differing in symptom severity (‘Low’ and ‘High’ are light and dark blue, respectively). (c) **Permutation test for differences in IS-RSA by symptom severity.** Vertical black line denotes z transformed difference (High-Low) in correlation coefficients in semantic-to-neural IS-RSA Colored histogram is a surrogate distribution comprised of randomly generated with shuffled severity labels. p value is derived non-parametrically (N = 25,000). Top left: Hippocampal patterns during ‘PTSD’ narratives. Top left: Hippocampal patterns during ‘PTSD’ narratives. Top right: PCC patterns during ‘PTSD’ narratives. Bottom left: Hippocampal patterns during ‘Sad narratives. Bottom right: PCC patterns during ‘Sad narratives (d) **Comparison of correlation coefficients across regions and symptom subgroups.** Bars reflect semantic-to-neural correlation coefficients (single values) for patterns in the hippocampus and PCC, shown for the ‘PTSD’ scripts across symptoms group (‘Low’: light red, ‘High’: dark red) and ‘Sad scripts (‘Low’: light blue, ‘High’: dark blue). Significance was assessed through permutation tests. HC – hippocampus, ** p < 0.01; *** p < 0.001, N.S – non-significant.

## Discussion

Despite continuous effort, the nature of intrusive traumatic autobiographical memories and the mechanisms underlying their unique perceptual attributes in PTSD remain largely unknown^47^. Here we used individualized traumatic autobiographical memory narratives in a script reactivation paradigm, in which PTSD patients listened to a novel rendition of their traumatic memory. We set out to ask whether, and how the hippocampus and amygdala differentiate traumatic autobiographical memories from sad ones. Given the duration and richness of the stimuli, our paradigm was at the intersection of autobiographical memory retrieval and naturalistic narrative comprehension tasks.

We leveraged the variance between idiosyncratic memories by quantifying their semantic similarity to ask whether their neural representations are altered during the processing of personal trauma narratives, compared to negative non-traumatic narratives of the same individuals (‘Sad’). Using intersubject representational similarity analysis, we found that hippocampal patterns showed a differentiation in semantic representation by narrative type; ‘Sad’ scripts which were semantically similar (e.g., death of a loved one) across participants, elicited similar neural representations. Conversely, thematically-similar traumatic autobiographical memories (of DSM- 5 Criterion A event) did not elicit similar representations. This effect was more pronounced in the left hippocampus and its posterior regions. Unlike the hippocampus, the amygdala did not represent semantic information in a significant manner, suggesting a poorer representational space for semantic content.

Semantic-to-neural mapping has been demonstrated comprehensively, both in the hippocampus^48, 49^ and cortex^50, 51^. Therefore, we expected to observe a link between narratives’ semantic similarity and elicited neural similarity. That said, here we make two advances: First – we extend this understanding into an underexplored domain: real-life traumatic autobiographical memories in PTSD. Second – we observe that within the same brains, hippocampal representations differed considerably between two types of autobiographical memory of comparable content and valence.

In a complementary approach, we decoded condition identity from the hippocampal patterns. The fact that we were able to tease the two negative conditions apart suggests that these signals hold some shared high-dimensional pattern implying a common cognitive state shared across participants.

Finally, we focused on the PCC to ask whether unlike the hippocampus, this region recently thought of as a cognitive bridge between the world events and representation of the self^45, 52^, will demonstrate a positive relation between semantic content and neural patterns of the traumatic narratives. We indeed observed such a relation in the PCC, with individual symptom severity mediating the extent of semantic-to-neural representation. Conversely, this differentiation by PTSD severity was not evident in the hippocampus.

Our key findings therefore are twofold: first – that the emotional content of autobiographical memories is represented differently in the two major systems subserving autobiographical memory - the hippocampus and the PCC. Second – that traumatic autobiographical memories undergo a parallel, or a dissociable mode of representation suggesting they profoundly differ from neurotypical autobiographical memories of comparable content and valence.

We also observed stronger semantic representation in the posterior subregion of the hippocampus, highlighting its function in the task’s demand for narrative imagery and memory retrieval of distant past^53^, rather than general PTSD symptomatology, in which the anterior hippocampus is generally assumed to take a greater role^54, 55^.

Why would traumatic autobiographical memories be represented differently than non-traumatic ones? We discuss several explanations; We speculate that dysfunctions in peritraumatic memory encoding, an impairment thought to lead to mnemonic sequelae collectively referred to as ‘memory fragmentation’^56, 57^ (but see ^58^ for an opposing view), may be the culprit of the poorer semantic representation of the traumatic memory. The weak semantic-to-neural mapping of traumatic narratives observed in our study resonates with reports of memory impairment attributed to the traumatic experience – memory disorganization, difficulty in voluntary retrieval and lack of narrative coherence^59, 60^.

We stress that although the scripts were directly portraying participants’ autobiographical memories, participants were naïve to these audio renditions. Basing off theories of memory fragmentation, it is possible that while for non-traumatic memories, script playback met some sort of a preexisting mental episodic structure of the events, in traumatic autobiographical memories they did not. In other words, to some extent, although extremely personal in nature, hippocampal signals imply that the traumatic narratives can be likened to non-personal stimuli. Another intriguing possibility is that patients attempted to block or suppress the reactivation of the traumatic content, and by doing so, exhibited brain activity that was incongruent with the semantic content presented to them^61, 62^. That said, we have no empirical way to test this alternative explanation. Re-examining the nature of these memories after successful trauma-focused psychotherapy may shed further light on the observed results.

The PCC (predominantly its left part) is known to be engaged during retrieval of autobiographical memory^63, 64^ and emotional narrative imagery^46^. Our data support these finding as well as the laterality difference. Moreover, two meta-analyses reported higher PCC activity during retrieval of autobiographical memory in PTSD compared to non-trauma exposed controls^64, 65^. The extent to which the PCC represented semantic content of the traumatic memory was dependent on individual symptom severity – the subgroup with elevated symptoms showed a closer relation between semantic content and neural signals. This was in contrast with hippocampal activity, where symptom severity had no explanatory power.

Copious reports tied alterations in PCC-hippocampus coupling to PTSD^66–68^, and demonstrated that individual changes in this link were associated with symptom severity^43, 69^. We speculate that both aforementioned explanations, namely lack of a preexisting narrative, and attempts at suppression, may have contributed to striking a balance in mentalization of the audio scripts by these two regions supporting autobiographical memory.

Finally, in light of the global alterations in brain structure and connectivity in PTSD, our within-subject design, where the traumatic narratives were compared to sad ones within the same individuals, controlled for these between-cohort effects.

Our study complements previous investigations of basic learning and memory mechanisms in PTSD by adding to our understanding of the processes that render a traumatic memory deviant from other non-traumatic memories. Moreover, we suggest that in addition, or in parallel, to the disturbance of fundamental learning and memory processing, high-level representation of meaning and content also undergo changes in PTSD.

In clinical settings, the evaluation of traumatic memory organization is often reliant mostly of meta-memory: the patient’s self-report about memory coherence of the traumatic experience ^58^. Semantic representation of idiosyncratic autobiographical memories using IS-RSA may allow a more objective neural marker for PTSD, and intriguingly, to amelioration of the disorder over the course of therapy. Importantly, to assign “coherence” to discrete memories, we must rely on the relational structure of semantic similarity generated from multiple memories of different individual. This way, semantic ‘anchors’ may be established, upon which an individual’s neural patterns can be evaluated.

We would like to acknowledge several limitations and shortcomings of our study: some semantic themes appeared more often in one narrative condition than the others (e.g., combat scenes were never regarded as a ‘Sad’ narrative). That said, we believe that the careful use of the same pool of sentences as well as the somatic induction, aided in rendering the scripts as comparable as possible. The IS-RSA did not uncover a significant representation of the ‘Calm’ narratives, which was unexpected since there is no reason to assume these kind of scripts were encoded in a different way. The variance in semantic content across ‘Calm’ narratives was much smaller than both ‘PTSD’ and ‘Sad’, and this may have obscured the relation to the neural patterns evoked by them. Given that positive autobiographical memories are more ubiquitous than traumatic ones, future studies should perhaps gently direct patients into recall of more diverse schemas during imagery development.

Ending with our initial question, the very nature of PTSD phenomenology remains an open question: is PTSD an extreme case of ‘standard’ negative emotional processing or a divergent cognitive entity altogether? Our main finding, that hippocampal patterns of PTSD patients showed a differentiation in semantic representation by narrative type during memory reactivation, supports the idea of a profoundly separate cognitive experience in the reactivation of traumatic memories.

## Methods

### Experimental paradigm

Twenty-eight participants (mean age = 38.1 ± 10.5 years, range = 24 – 63; females (n = 11), males (n = 17) took part in this study. All participants had chronic PTSD (see Table 1 for cohort demographics). PTSD diagnosis was established using the Clinician-Administered PTSD Scale (CAPS-5)^1^. The cohort in this study is part of a larger, longitudinal study focused on effects of ketamine-aided extinction in PTSD. Our data are based on the baseline session assessment, with no drugs being administered.

**Table 1:**
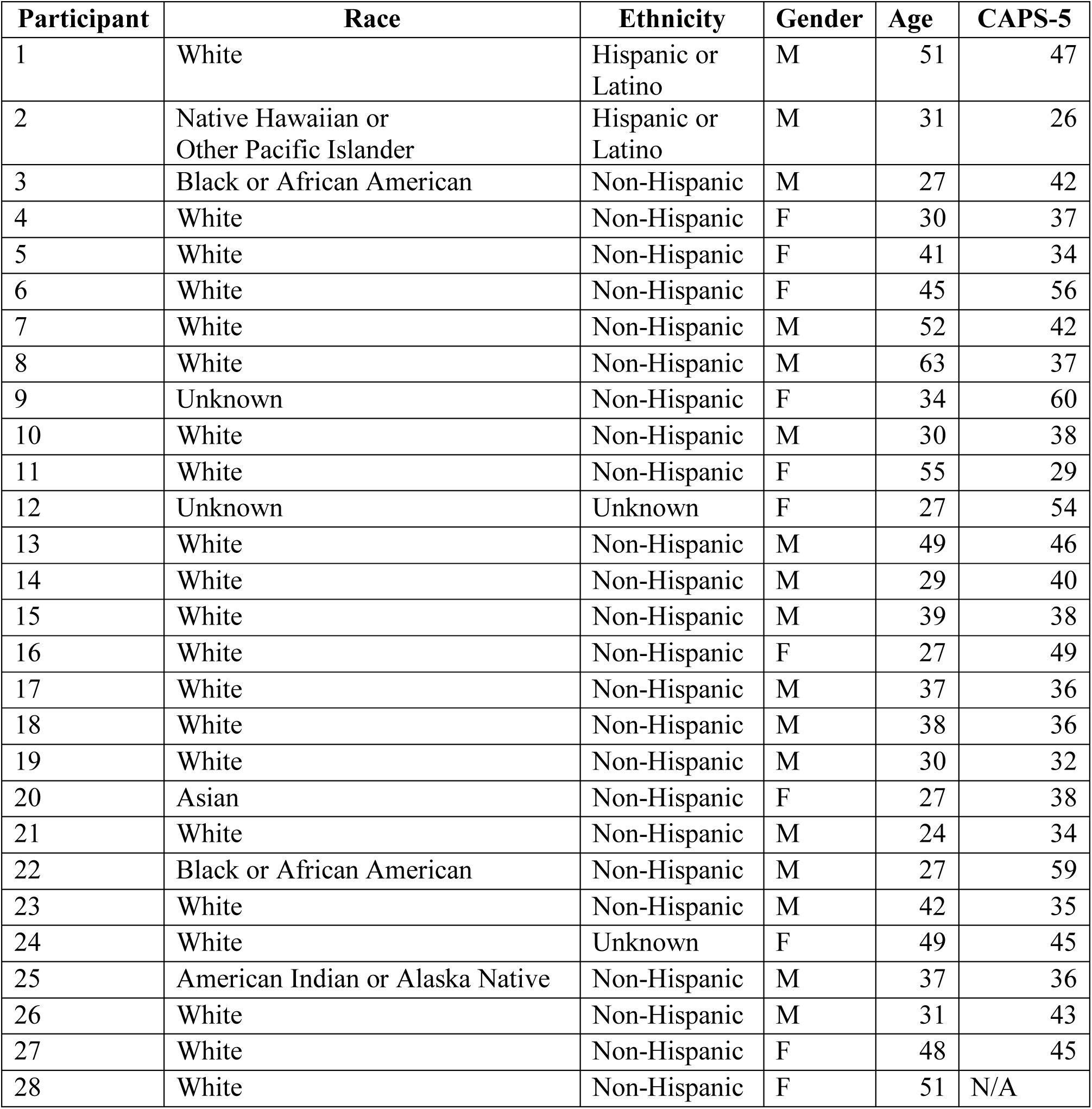
Cohort demographics

Exclusion criteria included a diagnostic history of bipolar disorder, borderline personality disorder, obsessive-compulsive disorder, schizophrenia or schizoaffective disorder, dementia, current psychotic features, or suicide risk, moderate or higher severity of substance use disorder and history of traumatic brain injury. Participants who were currently engaged in trauma focus therapy were also ineligible to participate in the study. Lastly, patients were excluded for acute medical illness. Psychotic features were determined by the Structured Clinical Interview for DSM-IV (SCID)^70^.

Participants completed an imagery-development procedure in which they were asked to describe the traumatic event associated with their PTSD, as well as a significant sad, but not traumatizing event (‘Sad’) and a positive, low-arousal, event in which they felt relaxed (referred to as ‘Calm’). The imagery scripting procedure followed a procedure presented by Sinha et al.^71^ Participants were asked to describe the events in as much detail as possible. They were then asked to select at least three physiological responses corresponding to each specific event to be later embedded in the narrative. Using this information, we developed audio scripts, approximately 120-second long for each event (Duration (sec): ‘PTSD’ = 120.19 ± 1.20; ‘Sad’ = 119.64 ± 1.48; ‘Calm’ = 118.69 ± 2.72), narrated by a male member of the research staff. The narratives were all in second-form pronouns (‘you’/’your’), mostly in the present tense.

The scripts were comprised of three main elements – the episodic unfolding of events (time and place, scenery description, actions, dialogues), description of mental state (e.g., ‘you feel helpless’, ‘a feeling of peace comes over you’), and a vocabulary of sensory and somatic phrases used to promote reexperiencing (e.g., ‘your heart beats faster’). The somatic vocabulary consisted of references to heartbeat, respiration, muscle tone, perspiration, tearing etc. Both negatively-valenced script types – ‘PTSD’ and ‘Sad’ – were conveyed using the same vocabulary, to control and maximize the similarity of reactivation between these conditions. The positive scripts (‘Calm’) often mentioned similar autonomic functions, but with opposite reactions (e.g., muscle relaxation, slow breathing).

### Semantic analysis

For the purpose of semantic analysis some names, for example places with greater geographical resolution than a US state, commercial brands or firms and specialized military jargon or acronyms were removed. Sentences were defined according to full stops, question, and exclamation marks as they appear in the text read by the narrator. As part of the reactivation procedure aimed to heighten autonomic arousal, the pace (words per script) of both negatively valenced scripts was intentionally higher and their duration slightly longer than the ‘Calm’ ones.

During preprocessing of the semantic input, punctuation marks were erased, texts were transformed to lower-case and tokenized. Stop words, defined by MATLAB R2020a’s default NLP vocabulary, were removed. Words underwent lemmatization and were then assigned a 300- dimension vector representation. The semantic space we used was a pre-trained embedding for 1M English words (16B tokens) available through MATLAB’s word2vec NLP tools. Vector representations were calculated for each single word. Words in the scripts which were not indexed in the pre-trained space were not included in the analysis. The semantic representation of the next level in text hierarchy – the sentence – were calculated as the average representation of words in it. Similarly, the semantic representation of each entire script was calculated by averaging the representation of its sentences.

Before computing similarity between scripts, we scaled the vector representations by subtracting the average norm of all words embedding in the entire vocabulary that was used in the study (based on all script types), following a normalization procedure described in^72^. The semantic similarity was derived from a transformed distance matrix of pairwise cosine distances of the vectorial representations. We used cosine distance as it is considered a better fit for semantic analysis than Euclidean distance because it does not consider vector magnitude, which is often biased in datasets involving text corpora due to differences in word occurrence^73, 74^.

### Dimensionality reduction

Dimensionality reduction of the semantic information was done using *t-*distributed stochastic neighbor embedding (*t*-SNE)^75^. Similarly to principal component analysis (PCA), *t-*SNE also reduces dimensionality of the input data set. It is superior however to PCA, in its ability to preserve local structure. Projection of *t-*SNE clusters to 3-dimensional space was carried out in MATLAB, with an intermediate step of a PCA into 100 components. Learning rate was set to 1000 and perplexity was set to 30.

### Experimental Procedure

All participants listened to each script type three times in the scanner. Participants were naïve to this new scripted rendition of their autobiographical memories and not familiar with the voice of the narrator. The order of scripts was fixed and identical across all participants but one. This order was chosen to prevent random occurrences of the same script consecutively. The order was: T*-*C-S-C-T-S-C-T-S in all scans but one, whose order was S-C-T-C-S-T-C-S-T (T = Trauma, S = Sad, C = calm). Each script began with a slide instructing the participant to press a button to initiate the next playback and its type. No visual information was displayed during script playback.

### MRI Scan

MRI data were collected with a Siemens 3T Prisma scanner, using a 32-channel receiver array head coil. High-resolution structural images were acquired by Magnetization-Prepared Rapid Gradient-Echo (MPRAGE) imaging (TΡ = 1 s, TE = 2.77 ms, TI = 900 ms, flip angle = 9°, 176 sagittal slices, voxel size = 1 ×1 × 1 mm, 256 × 256 matrix in a 256 mm FOV). Functional MRI scans were acquired while the participants were listening to the narrated scripts, using a multi- band Echo-Planar Imaging (EPI) sequence (multi-band factor =4, TR= 1000 ms, TE= 30 ms, flip angle = 60°, voxel size = 2 × 2× 2 mm, 60 2 mm-thick slices, in-plane resolution = 2 × 2 mm, FOV = 220 mm).

### MRI preprocessing

Data were preprocessed with fMRIPrep, version 20.2.0^76^. For a complete preprocessing procedure please refer to Supplementary Information. Functional images were motion- and slice-time corrected, aligned to T1 anatomical images, and then warped to MNI space. Analysis of the functional data included the following regressors: 6 movement variables (translation and rotation), framewise displacement, the first 6 anatomical components based noise correction (CompCor) and the 6 first discrete cosine regressors. Subsequent preprocessing and statistical contrasts were done using standard statistical parametric mapping (SPM12, Wellcome Department of Imaging Neuroscience) algorithms (fil.ion.ucl.ac.uk/spm) and custom MATLAB R2018a code.

### ROI analysis

Given our a-priori interest in the function of amygdala and hippocampus in PTSD, we defined masks for ROI analysis of these structures, bilaterally using the probabilistic Harvard Oxford atlas ^77^ thresholded at 25%^78^. Prior to ROI analysis, functional images underwent spatial smoothing using a Gaussian kernel of 1 mm full-width at half maximum (FWHM) to enhance signal-to-noise ratio and classification accuracy^79^. Time courses were extracted from the entire session and were applied with a discrete cosine transform high-pass filter (cutoff of 128 sec)^80^. ROI data were then normalized using z-score. Spikes in the data, exceeding 4 times the voxel’s standard deviation were applied de-spiking and were interpolated using the mean of one TRs engulfing each side of the outlier data point. In line with previous studies, functional data were shifted 5 seconds (5 TRs) to account for the delay in the hemodynamic response compared with the audio stimuli^72, 81, 82^. The segments containing the script narratives were extracted with time indices rounded to include the nearest TR interval to prevent omission of functional data.

Each script was played three times during the scan. To enhance signal to noise in identifying the recurring pattern elicited by the script, the time course for each ROI was averaged across the three repeats of each script. The voxel time course of this average run was collapsed across time to generate a spatial pattern associated with the reactivation of the specific autobiographical memory. This approach also aided in circumventing the slight mismatches in script durations across participants that would have been detrimental for similarity based on temporal fluctuations.

### Intersubject representational similarity analysis (IS-RSA)

To relate the semantic content with neural representation and to determine whether this representation differs in PTSD-related narratives, we conducted an intersubject representational similarity analysis (IS-RSA)^73, 83–85^. The neural similarity matrix was derived from each ROI separately. The semantic similarity matrix was fixed in all analyses (for some analyses it was broken down to smaller matrices, for example, sub-cohorts differing in symptoms severity).

A neural similarity matrix was then generated by calculating pair-wise Pearson correlation for each pair of the 84 scripts. Metrics that are represented as distance by default (rather than similarity) were transformed to similarity using

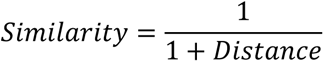

Since this study comprised of 28 participants, who each listened to 3 different scripts, the narrative-based similarity matrices (e.g., semantic, acoustic) consisted of the lower triangle of a 28 X 28 matrix, extracted separately per narrative type (‘PTSD’, ‘Sad’, ‘Calm’). This corresponded to 378 unique combinations when vectorized ((28 X (28 – 1)) / 2 = 378). Similarly, trait-based similarity matrices (e.g., CAPS) consisted of the lower triangle of a 28 X 28 matrix, corresponding to 378 unique combinations. Of note, in our design both the neural responses and the naturalistic stimuli varied across individuals.

We tested the significance of IS-RSA by subjecting p-values of Spearman’s correlation coefficients amassed across ROIs and conditions (e.g., 2 ROIs X 3 script types) by using a false-discovery rate (FDR) at q = 0.05 implemented in MATLAB R2018a as function ‘fdr_bh’.

To compare correlation coefficients between narrative types (e.g., ‘PTSD’ vs. ‘Sad’) we used two methods depending on groups’ dependency: In cases where one variable is shared (e.g., correlation between CAPS and neural similarity in ‘PTSD’ narratives compared with correlation between the same CAPS data and neural similarity in ‘Sad’ narratives) we used Steiger test^86^ as implanted in ‘r_test_paired.m’ in MATLAB. Briefly, each correlation coefficient is converted into a z-score using Fisher’s r-to-z transformation. Next the asymptotic covariance of the estimates is computed and are then used in an asymptotic z-test. We reported p-values from a two-tailed probability distribution. In contrast, in cases where the two comparisons had no shared components (e.g., correlation between semantic and neural similarity in ‘PTSD’ narratives and correlation between semantic and neural similarity in ‘Sad’ narratives), we used the ‘corr_rtest’ function in MATLAB to convert both correlation coefficient into z-score using Fisher’s r-to-z transformation and calculate their absolute difference. This value was assigned a p value from a cumulative normal distribution function (‘normcdf’ in MATLAB). We report p-values from a two-tailed probability distribution.

### IS-RSA visualization

IS-RSA is usually calculated using nonparametric tests (Spearman, Kendall). However, the limitation of Spearman, being a ranked test, is that it does not yield a regression in the same manner linear correlations do. Due to this technical limitations in visualizing the relationship between data matrices in the IS-RSA, we plotted slopes that are computed based on Pearson correlation. Note that despite this difference in visualization, throughout this study, the correlation coefficient reported and discussed are Spearman’s ρ values.

### Longitudinal parcellation of the hippocampus

To date, several definitions for segmentation of long axis of the human hippocampus exists. We followed a percentile-based segmentation of the hippocampus into three regions along its long axis, described by Poppenk et al.^41^. To avoid discrepancies between boundaries defined by the various segmentation methods, we omitted the medial part from our analyses and focused only on the two extremities – the most anterior and posterior thirds.

### Neural general linear modeling (GLM) and contrasts

We conducted general linear modeling (GLM) of the functional scans of each participant by modeling the observed BOLD signals and regressors to identify the relationship between the task events and the hemodynamic response. First, functional data underwent spatial smoothing using a 6 mm FWHM kernel (note that a different kernel was used in IS-RSA). Next, regressors related to all events were created by convolving a train of delta functions representing the sequence of individual events with the default basis function in SPM12, which consists of a synthetic hemodynamic response function composed of two gamma functions. The GLM included three separate regressors for the onset of the three types of scripts – ‘PTSD’, ‘Sad’ and ‘Calm’. We carried out linear contrasts of parameter estimates to identify effects in each participant. Statistical maps from all participants were then entered into a second-level group analysis to implement a random-effects statistical model.

### Classification analysis

We decoded narrative conditions (‘PTSD’ or ‘Sad’ script) from multivoxel spatial patterns data using a rLDA (regularized linear discriminant analysis classifier, ‘fitdiscr’ function in MATLAB) which shows superior performance over LDA in high-dimensional imaging data which may present issues of multicollinearity and overfitting. Data from all script repeats of the two negatively-valenced conditions, along with corresponding condition labels, were used to train the rLDA. Input data consisted of 28 participants X 2 conditions X 3 repeats yielding a 168-samples in total. To test the model’s performance on a testing data set we iteratively repeated this process with permuted data partitions (N = 2,500) per ROI and condition for cross validation and then applied cross-validation.

### Acoustic similarity analysis

Acoustic similarity of the scripts was computed based on acoustic landmarks envelope. We used a custom MATLAB script, adapted from Oganian et al.^87^ to extract the analytic envelope of the speech signal filtered within critical bands based on the Bark scale which is a psychoacoustical measure of loudness.

### Whole-brain investigation

The main contrast used in the whole-brain investigation was “All Scripts > Baseline” where all script event types (i.e., ‘PTSD’, ‘Sad’ and ‘Calm’) were contrasted with the baseline interval between scripts. Statistical inference was made based on whole-brain statistical maps corrected for multiple comparisons using cluster size threshold family-wise error rate of p_(FWE)_ < 0.01 for the identification and extraction of regions of interest (ROIs).

### Symptom severity analysis

CAPS - PTSD diagnosis was established using the Clinician-Administered PTSD Scale (CAPS-5)^1^. CAPS was administered within one month of the imaging session. The questionnaire data of one participant were missing and were interpolated using the group’s mean per questionnaire item.

## Code availability

The scripts used for data analysis are available in https://osf.io/dc7jb/

fMRI preprocessing was done in fMRIPrep, analyses were conducted primarily in MATLAB R2018b and R2020a (MathWorks, Natick, MA)

## Data availability

Data supporting the findings of this study are deposited in https://osf.io/dc7jb/

## Acknowledgments

We thank Drs. Temidayo Orederu and Yaara Yeshurun for fruitful discussions. We also thank Drs. Aya Ben Yakov and Aharon Ravia for their advice on analysis and preprocessing.

## Ethics declarations

The authors declare no competing financial interests.

---

**Supplementary Figure 1:**
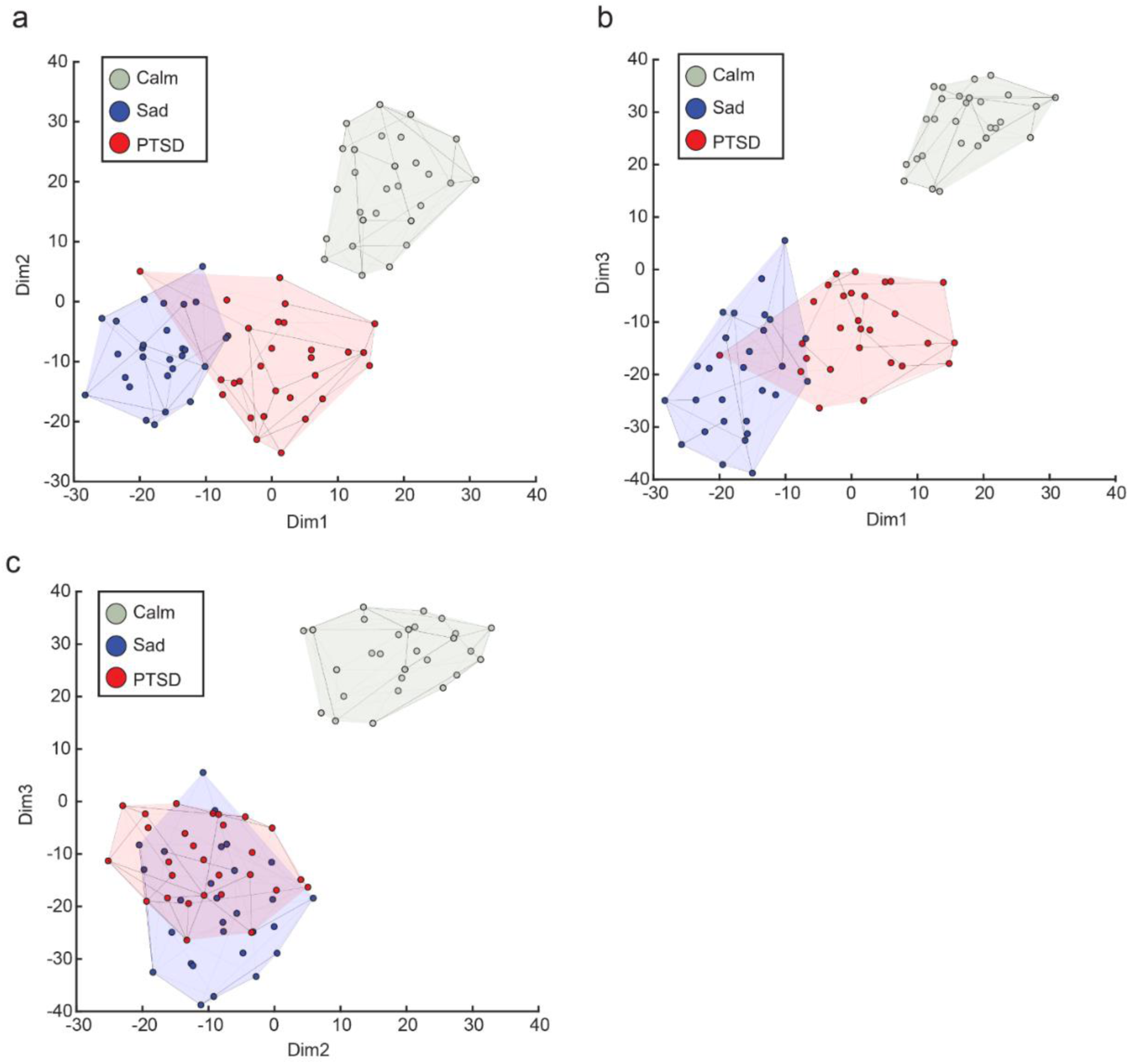
***(a)* projections of semantic space based on three principal dimensions**. t-SNE embedding of scripts projected onto dimensions 1-2. Each dot represents a single script, Colored volumes are continuous spaces occupied by each script type. Color denotes script type (‘PTSD’: red, ‘Sad’: blue, ‘Calm’: gray). Note overlap of ‘PTSD’ and ‘Sad’ semantic content. ***(b)*** Same as **(a)** but projected onto dimensions 1-3. ***(c)*** Same as **(a)** but projected onto dimensions 2-3.

**Supplementary Figure 2:**
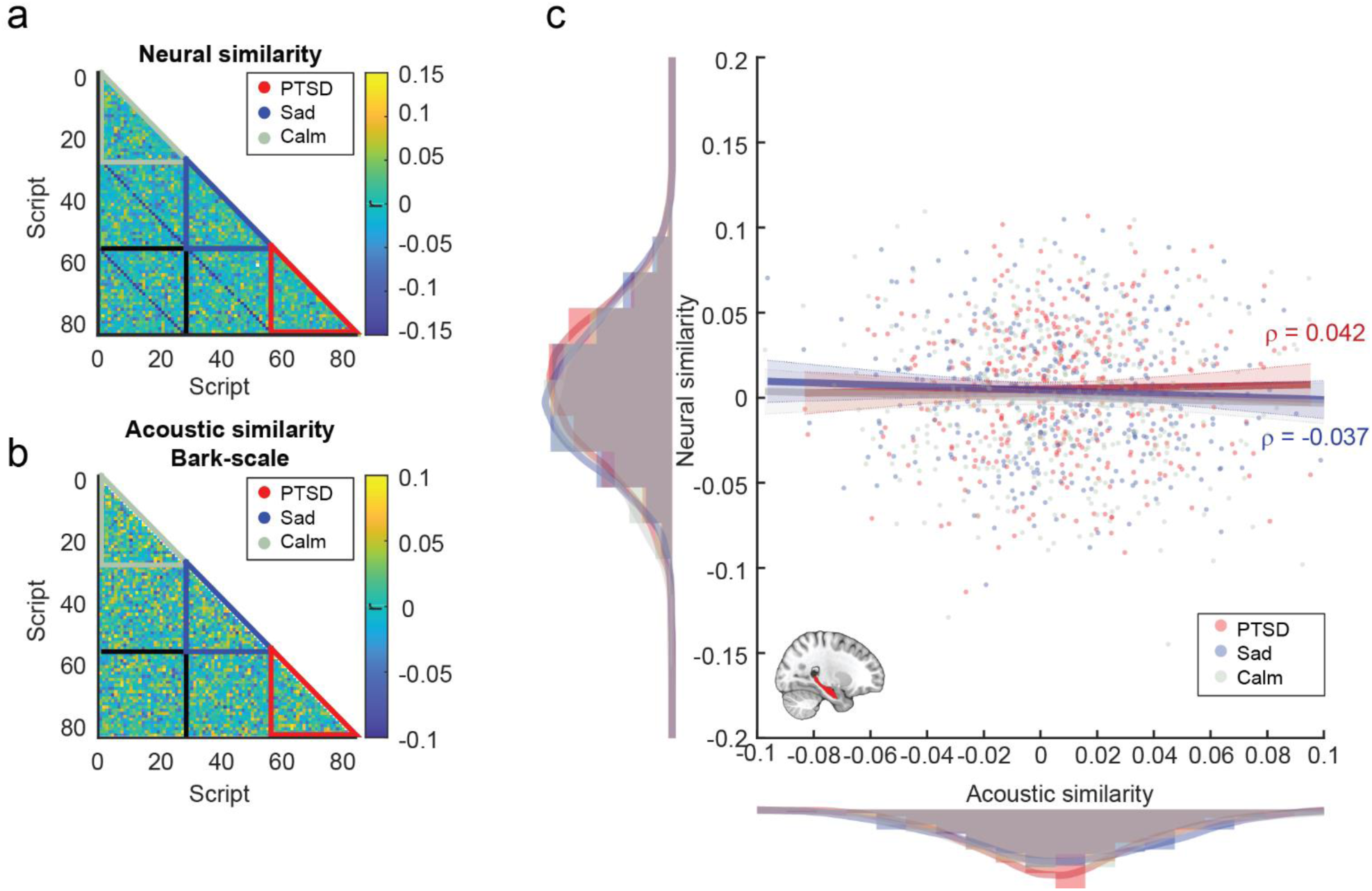
***(a)* Hippocampus – neural similarity matrix.** Script-by-script neural similarity matrix for spatial patterns extracted from the hippocampus during script reactivation. Within-category similarity is marked in colored triangles off the matrix diagonal (‘PTSD’: red, ‘Sad’: blue, ‘Calm’: gray). ***(b)* Acoustic similarity matrix.** Acoustic similarity of scripted narratives. Within-category similarity is marked in colored triangles off the matrix diagonal (‘PTSD’: red, ‘Sad’: blue, ‘Calm’: gray). ***(c)* Hippocampus – acoustic-to-neural IS-RSA.** Intersubject representational similarity analysis conducted on pairwise similarity of acoustic and neural patterns in the hippocampus. Each datapoint is one pairwise comparison. Analysis was iterated per script type (‘PTSD’: red, ‘Sad’: blue, ‘Calm’: gray). Histograms along axes depict similarity distribution, thick trace depict estimated density, colors correspond to main legend. Regression lines are approximate visualization of Spearman correlation rho coefficients for IS-RSA in ‘PTSD’ and ‘Sad’ scripts (red and blue resp.).

**Supplementary Figure 3:**
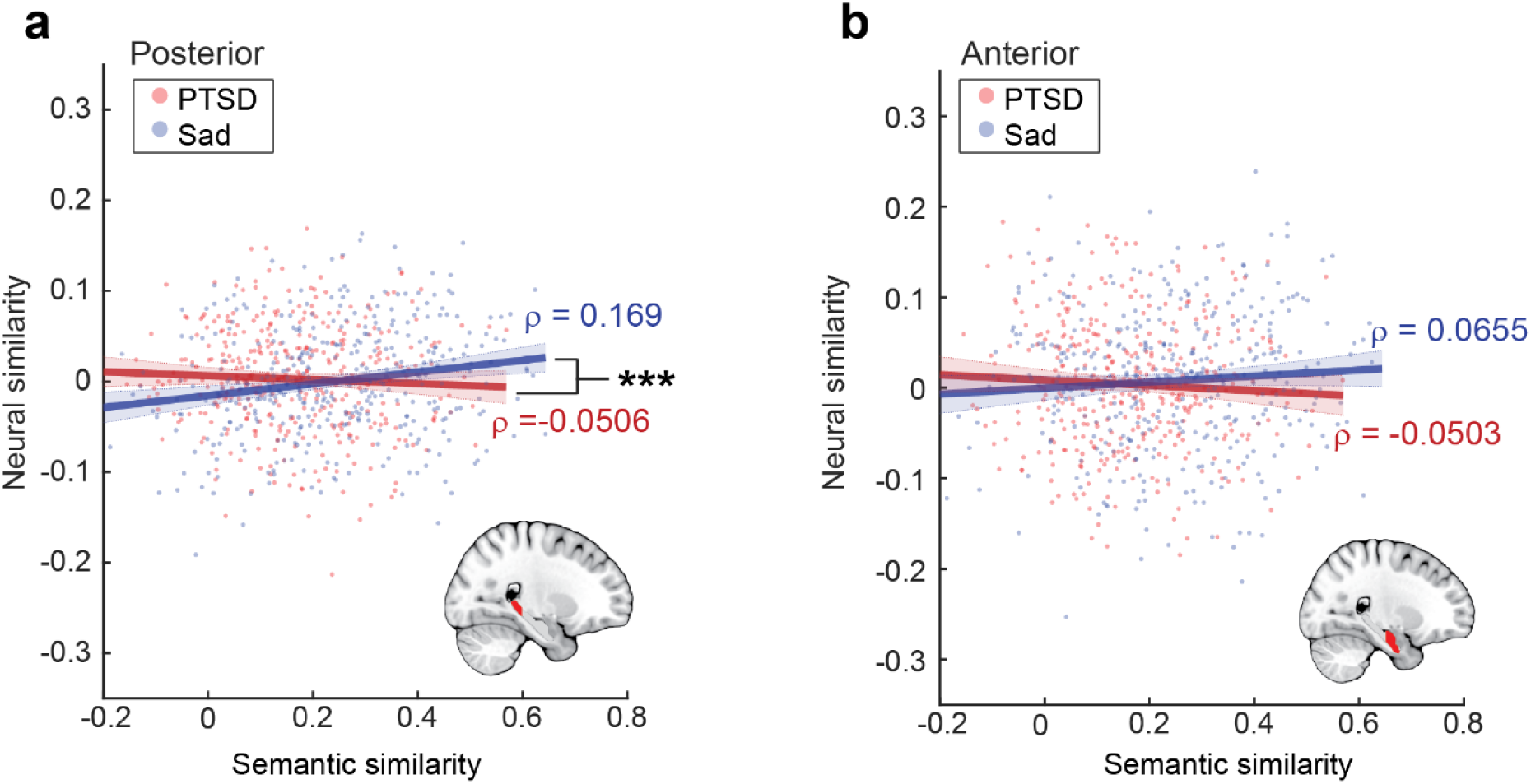
*(a)* **semantic-to-neural IS-RSA in posterior hippocampus.** Intersubject representational similarity analysis conducted on pairwise similarity of semantic content and neural patterns in the posterior hippocampus. Each datapoint is one pairwise comparison. Analysis was iterated per script type (‘PTSD’: red, ‘Sad’: blue). Regression lines are approximate visualization of Spearman correlation rho coefficients for IS-RSA in ‘PTSD’ and ‘Sad’ scripts (red and blue resp.). * p < 0.001. *(b)* Same as **(a)** but for the anterior hippocampus.

**Supplementary Figure 4:**
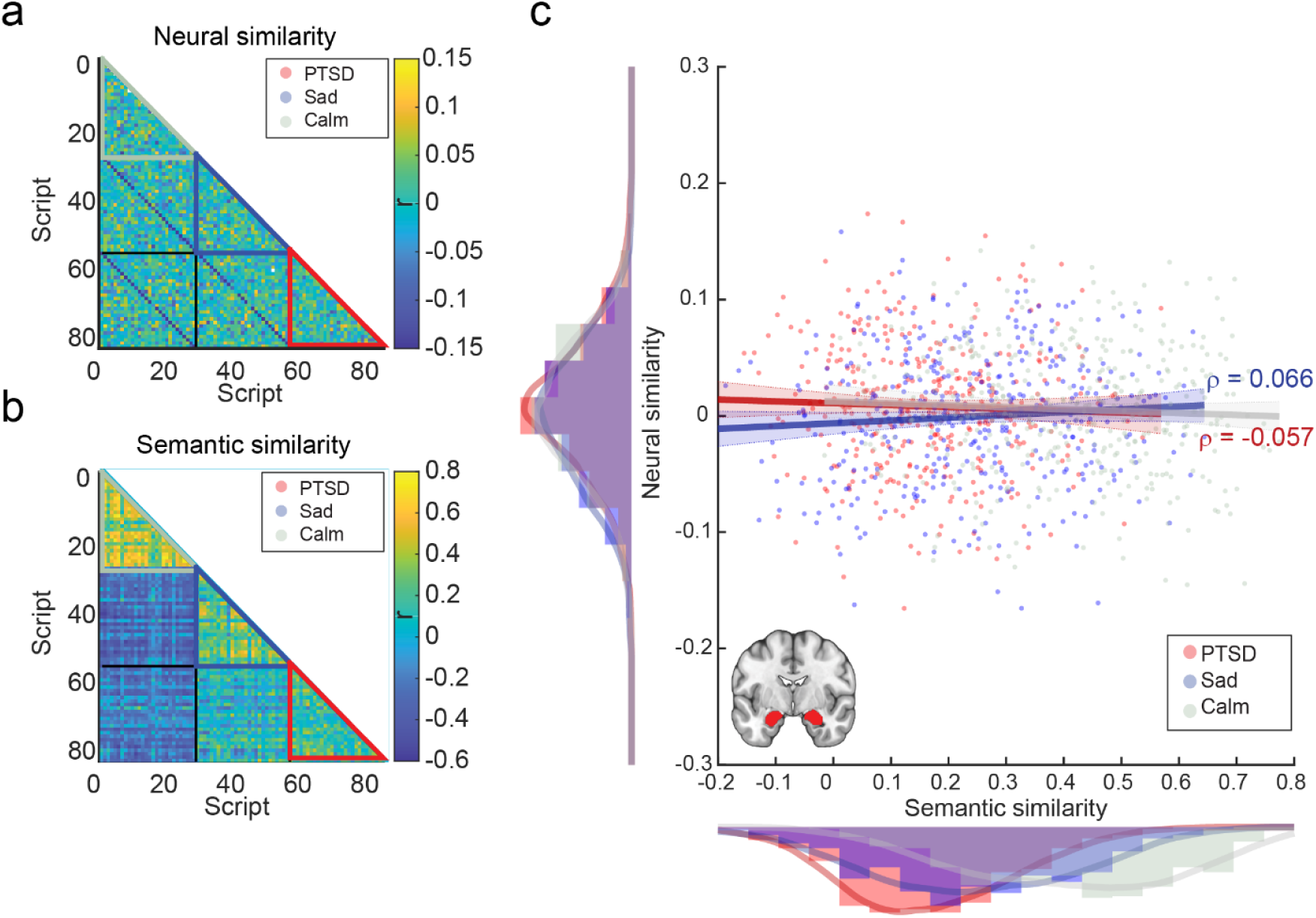
**(a) Amygdala – neural similarity matrix.** Script-by-script neural similarity matrix for spatial patterns extracted from the amygdala during script reactivation. Within-category similarity is marked in colored triangles off the matrix diagonal (‘PTSD’: red, ‘Sad’: blue, ‘Calm’: gray). **(b) Semantic similarity matrix.** Semantic similarity (Pearson’s correlation coefficient, r) of scripted narratives. Within-category similarity is marked in colored triangles off the matrix diagonal (‘PTSD’: red, ‘Sad’: blue, ‘Calm’: gray). **(c) Amygdala – semantic-to-neural IS-RSA.** Intersubject representational similarity analysis conducted on pairwise similarity of semantic content and neural patterns in the amygdala. Each datapoint is one pairwise comparison. Analysis was iterated per script type (‘PTSD’: red, ‘Sad’: blue, ‘Calm’: gray). Histograms along axes depict similarity distribution, thick trace depict estimated density, colors correspond to main legend. Regression lines are approximate visualization of Spearman correlation rho coefficients for IS-RSA in ‘PTSD’ and ‘Sad’ scripts (red and blue resp.).

## Supplementary methods

### fMRIprep Preprocessing

Results included in this manuscript come from preprocessing performed using fMRIPrep 20.2.0 (Esteban, Markiewicz, et al. (2018); Esteban, Blair, et al. (2018); RRID:SCR_016216), which is based on Nipype 1.5.1 (Gorgolewski et al. (2011); Gorgolewski et al. (2018); RRID:SCR_002502).

### Anatomical data preprocessing

A total of 1 T1-weighted (T1w) images were found within the input BIDS dataset. The T1- weighted (T1w) image was corrected for intensity non-uniformity (INU) with N4BiasFieldCorrection (Tustison et al. 2010), distributed with ANTs 2.3.3 (Avants et al. 2008, RRID:SCR_004757), and used as T1w-reference throughout the workflow. The T1w-reference was then skull-stripped with a Nipype implementation of the antsBrainExtraction.sh workflow (from ANTs), using OASIS30ANTs as target template. Brain tissue segmentation of cerebrospinal fluid (CSF), white-matter (WM) and gray-matter (GM) was performed on the brain-extracted T1w using fast (FSL 5.0.9, RRID:SCR_002823, Zhang, Brady, and Smith 2001). Volume-based spatial normalization to one standard space (MNI152NLin2009cAsym) was performed through nonlinear registration with antsRegistration (ANTs 2.3.3), using brain-extracted versions of both T1w reference and the T1w template. The following template was selected for spatial normalization: ICBM 152 Nonlinear Asymmetrical template version 2009c [Fonov et al. (2009), RRID:SCR_008796; TemplateFlow ID: MNI152NLin2009cAsym],

### Functional data preprocessing

For each of the 1 BOLD runs found per subject (across all tasks and sessions), the following preprocessing was performed. First, a reference volume and its skull-stripped version were generated using a custom methodology of fMRIPrep. Susceptibility distortion correction (SDC) was omitted. The BOLD reference was then co-registered to the T1w reference using flirt (FSL 5.0.9, Jenkinson and Smith 2001) with the boundary-based registration (Greve and Fischl 2009) cost-function. Co-registration was configured with nine degrees of freedom to account for distortions remaining in the BOLD reference. Head-motion parameters with respect to the BOLD reference (transformation matrices, and six corresponding rotation and translation parameters) are estimated before any spatiotemporal filtering using mcflirt (FSL 5.0.9, Jenkinson et al. 2002). BOLD runs were slice-time corrected using 3dTshift from AFNI 20160207 (Cox and Hyde 1997, RRID:SCR_005927). The BOLD time-series (including slice-timing correction when applied) were resampled onto their original, native space by applying the transforms to correct for head-motion. These resampled BOLD time-series will be referred to as preprocessed BOLD in original space, or just preprocessed BOLD. The BOLD time-series were resampled into standard space, generating a preprocessed BOLD run in MNI152NLin2009cAsym space. First, a reference volume and its skull-stripped version were generated using a custom methodology of fMRIPrep. Several confounding time-series were calculated based on the preprocessed BOLD: framewise displacement (FD), DVARS and three region-wise global signals. FD was computed using two formulations following Power (absolute sum of relative motions, Power et al. (2014)) and Jenkinson (relative root mean square displacement between affines, Jenkinson et al. (2002)). FD and DVARS are calculated for each functional run, both using their implementations in Nipype (following the definitions by Power et al. 2014). The three global signals are extracted within the CSF, the WM, and the whole-brain masks. Additionally, a set of physiological regressors were extracted to allow for component-based noise correction (CompCor, Behzadi et al. 2007). Principal components are estimated after high-pass filtering the preprocessed BOLD time-series (using a discrete cosine filter with 128s cut-off) for the two CompCor variants: temporal (tCompCor) and anatomical (aCompCor). tCompCor components are then calculated from the top 2% variable voxels within the brain mask. For aCompCor, three probabilistic masks (CSF, WM and combined CSF+WM) are generated in anatomical space. The implementation differs from that of Behzadi et al. in that instead of eroding the masks by 2 pixels on BOLD space, the aCompCor masks are subtracted a mask of pixels that likely contain a volume fraction of GM. This mask is obtained by thresholding the corresponding partial volume map at 0.05, and it ensures components are not extracted from voxels containing a minimal fraction of GM. Finally, these masks are resampled into BOLD space and binarized by thresholding at 0.99 (as in the original implementation). Components are also calculated separately within the WM and CSF masks. For each CompCor decomposition, the k components with the largest singular values are retained, such that the retained components’ time series are sufficient to explain 50 percent of variance across the nuisance mask (CSF, WM, combined, or temporal). The remaining components are dropped from consideration. The head-motion estimates calculated in the correction step were also placed within the corresponding confounds file. The confound time series derived from head motion estimates and global signals were expanded with the inclusion of temporal derivatives and quadratic terms for each (Satterthwaite et al. 2013). Frames that exceeded a threshold of 0.5 mm FD or 1.5 standardized DVARS were annotated as motion outliers. All resamplings can be performed with a single interpolation step by composing all the pertinent transformations (i.e., head-motion transform matrices, susceptibility distortion correction when available, and co-registrations to anatomical and output spaces). Gridded (volumetric) resamplings were performed using antsApplyTransforms (ANTs), configured with Lanczos interpolation to minimize the smoothing effects of other kernels (Lanczos 1964). Non-gridded (surface) resamplings were performed using mri_vol2surf (FreeSurfer).

Many internal operations of fMRIPrep use Nilearn 0.6.2 (Abraham et al. 2014, RRID:SCR_001362), mostly within the functional processing workflow. For more details of the pipeline, see the section corresponding to workflows in fMRIPrep’s documentation.

### Copyright Waiver

The above boilerplate text was automatically generated by fMRIPrep with the express intention that users should copy and paste this text into their manuscripts unchanged. It is released under the CC0 license.

